# Elevated CO_2_ induces phyllosphere community changes in soybean

**DOI:** 10.64898/2025.12.08.693023

**Authors:** Connor Morozumi, Elizabeth A. Ainsworth, Katy D. Heath, Natalie Christian

**Author notes:** Email addresses. Author contributions: NC, KH: conceptualization; NC: methodology; CM and NC: formal analysis; CM and NC: investigation; LA: resources; CM: data curation; CM, and NC: writing - original draft; CM, NC, LA, KH: writing - review & editing; CM: visualization; NC: supervision; NC, KH, LA: funding acquisition.

## Abstract

Plant-associated microbiomes play significant roles in determining their hosts’ responses to stress, which is increasingly common in a rapidly changing world. For example, foliar endophytes can increase tolerance to drought and decrease herbivory. Global atmospheric changes, such as increased atmospheric carbon dioxide concentration ([CO_2_]), have the potential to directly and indirectly alter microbiome community structure in ways that affect host plants underscoring our need to understand how microbial communities *in planta* will change under global change. Here, we used a metabarcoding approach to assess community composition and network structure of bacterial and fungal phyllosphere microbiomes within soybean (*Glycine max*) exposed to ambient and elevated [CO_2_] in the field. We found that fungal community composition differed between soybean grown in elevated and ambient [CO_2_], while bacterial communities did not. Additionally, co-occurrence networks for both fungi and bacteria in elevated [CO_2_] exhibited marked changes compared to the ambient networks including the composition of hub taxa. Overall, our findings suggest that anthropogenic change, such as elevated [CO_2_], can cause profound shifts in community assembly of microbes within their plant hosts. Our findings may aid those developing agro-ecological strategies using microbes to improve crop traits, which necessitates understanding the drivers of microbiome community structure.

**Highlight:** Leaf-associated microbes benefit plants, but how climate change affects these communities remains unclear. We found that elevated CO_2_ shifted fungal and bacterial community composition and network structure within soybean leaves.

## Introduction

The present era is characterized by rapid, global climate change that is currently driving massive ecological and evolutionary changes in organisms across the planet (Intergovernmental Panel on Climate Change (IPCC), 2023). These changes may have similarly large effects on microbial communities. For microbes which are constituents of plant microbiomes, these resultant community-level shifts can cascade to affect plant health. For example, within various plant hosts and ecosystems, microbes are known to play vital roles in plant growth (Trivedi *et al*., 2020), crop yield (Li *et al*., 2022), disease resistance (Van Wees *et al*., 2008), and drought tolerance (Ma *et al*., 2020). Although we know that microbial communities are sensitive to abiotic conditions, and that changes in abiotic stress have downstream effects on hosts, the precise nature of how microbial communities shift in response to global climate change remains a critical outstanding question. This is particularly true for leaf microbiomes, whose responses to global change factors remain relatively unexplored compared to root microbiomes, despite the strong benefits that foliar microbes confer to plant health such as pathogen protection (e.g., Christian *et al*., 2017) and drought resistance (e.g., Giauque *et al*., 2019). Because leaf microbiota play key roles in mediating host responses to stress, we need to better understand how these microbial partners—as well as their interactions—are shifting under global change.

Elevated carbon dioxide concentration ([CO_2_]) in the atmosphere alters microbiomes within plants. Research examining belowground microbiomes demonstrated that elevated [CO_2_] often shifts microbial communities in soil (Wang *et al*., 2017; Usyskin-Tonne *et al*., 2020), the rhizosphere (Yu *et al*., 2018), and roots (Usyskin-Tonne *et al*., 2020). Collectively, this work has underscored that anthropogenic change may adversely affect plant microbial communities.

However, much research to date has focused on microbiomes of soil and the rhizosphere, with leaf microbiomes chronically understudied compared to belowground compartments. In the handful of aboveground studies to date, research has found elevated [CO_2_] shifts leaf microbial communities, though the effects of elevated [CO_2_] are dependent on tissue type and seasonal conditions (Bakker *et al*., 2023) as well as nitrogen fertilization rates and plant development stage (Ren *et al*., 2015). Elevated [CO_2_] may increase available niche space via increases to aboveground plant biomass and carbohydrate assimilation yet it also is known to decrease rates of stomatal opening (Ainsworth *et al*., 2002), a main route of transmission for internal leaf colonizers such as foliar endophytes (Huang *et al*., 2018).

Global changes such as increasing [CO_2_] are likely to affect not only the composition of microbiome communities, but also the interactions within these communities. It is complicated to study species interactions for species-rich microbial communities because of the sheer number of interactions present as well as the inherent difficulty of directly observing such interactions. Co-occurrence networks, while not directly interpretable as evidence for species interactions, have been employed as a way to depict and represent microbial communities (Layeghifard *et al*., 2017). For example, networks have been used to identify roles certain microbes appear to play in the microbiome. This is mainly through analyses such as identifying highly connected taxa (i.e., taxa with high centrality) or those that appear to play keystone roles in the network such as those acting as a “hub” (Agler *et al*., 2016). Examining the strength and direction of co-occurrences can also aid in understanding how taxa are potentially promoting or alternatively competitively excluding / partitioning niches in relation to one another. This is important given that global change is predicted to alter species interactions including predictions to either shift interactions away from mutualistic in nature (Tylianakis *et al*., 2008) or conversely might tighten positive relationships (He *et al*., 2013). Networks may detect meaningful, functional changes in community structure and interactions that might be obscured in other types of community-level analyses. In this way, network analyses have elucidated microbial interactions between suites of microbes and their hosts (e.g., Poudel *et al*., 2016) and have helped determine assembly rules for microbes (e.g., Liu *et al*., 2019), which could become increasingly important in a changing world rife with artificial bottlenecks, movement of species, and extinctions that reshuffle and reassemble communities.

Soybean (*Glycine max*) is an ideal system in which to study the effects of climate change on leaf microbiomes and community interactions. Soybean is a globally important crop due to its high protein and nutritional quality. In the United States, soybean is the second-most-planted field crop and the dominant oilseed crop, and its physiological responses to climate change have been extensively documented (Ainsworth *et al*., 2002; Ainsworth and Long, 2021). Soybean is an N-fixing species well-recognized for its belowground symbiosis with rhizobia bacteria, but like all plant species it also harbors a diverse microbiome. These communities of microbes differ by tissue type (Longley *et al*., 2020) and appear to be assembled via both vertical (Moroenyane *et al*., 2021) and horizontal (Liu *et al*., 2019) transmission routes. Most past research has been on rhizospheric and root nodule communities, with an emphasis on bacterial diversity.

Contrastingly, aboveground microbiomes of soybean are far less characterized or understood. In the few previous studies on aboveground organs, foliar communities of microbes in soybean appear to be driven by environmental (Christian *et al*., 2021) and host genotype effects (Zhou *et al*., 2023) as well as via temporal turnover across the season (Copeland *et al*., 2015). These foliar endophytes can alter plant gene expression and improve plant health in a context-dependent manner (Sosa Marquez *et al*., 2025). In a previous study using culture-based approaches to test foliar microbial species interactions under experimental global atmospheric change, researchers found that elevated [CO_2_] changed the abundance and competitive dynamics of a *Methylobacterium sp.* that was an antagonist to key *Colletotrichum* strains (Christian *et al*., 2021). This work was *in vitro* and looked only at pairwise interactions and no previous work has taken a metabarcoding or network approach to further contextualize and understand how these key microbes are structured into greater communities.

To understand how the phyllosphere microbiome within soybean responds to increased [CO_2_], we used a culture-independent metabarcoding approach to assess community and network structure of bacterial and fungal endophytic communities in soybean leaves grown under elevated and ambient [CO_2_]. We implemented a Free Air CO_2_ Enrichment (FACE) experiment to manipulate [CO_2_] in the field without enclosures, allowing plants to be grown under higher concentrations of CO_2_ without changing other abiotic factors such as air temperature and humidity (Ainsworth and Long, 2021). Additionally, these free-air systems allow for environmental transmission of endophytes. We hypothesized that leaf endophyte community composition—including both fungal and bacterial taxa—would differ between soybean grown in ambient and elevated [CO_2_], as has been shown in other plant compartments in previous FACE experiments (Wang *et al*., 2017). We expected these changes would largely be driven by changes to niche space (i.e., increased carbohydrates in leaf tissues under elevated [CO_2_], (Ainsworth *et al*., 2002)) and alterations to stomata which are key routes of endophyte immigration into leaf tissues. Previous studies indicate that taxonomic groups respond differently (Yu *et al*., 2016, 2018; Wang *et al*., 2017), but due to the variability of past studies we did not have *a priori* predictions of direction and/or magnitudes of these changes. Additionally, we hypothesized that elevated [CO_2_] would provide more photosynthates to leaf endophytes ultimately leading to increased competition within co-occurrence networks. Thus we predicted networks would have fewer links yet the remaining links would be more negative in nature given that 1) elevated [CO_2_] appears to benefit some taxa preferentially that are known to interact competitively (Christian *et al*., 2021) and 2) that mutualistic interactions in general are predicted to fare poorly under future global change (Tylianakis *et al*., 2008).

### Experimental Procedures

#### Field site, soybean cultivation and FACE experimental design

We conducted this study in a 16-ha field of soybean (*Glycine max*) at the SoyFACE facility located at the University of Illinois Urbana-Champaign (40 ° 02’N, 88 ° 14’W; https://soyface.illinois.edu). Soybean cultivar Pioneer P31T44E (Pioneer, Johnston, IA) was planted on 1 June, 2020 in 38.1-cm rows and planting density of ca. 445,000 plants per hectare. Experimentation infrastructure for CO_2_ enrichment was installed directly after planting. Plants were blocked into four experimental units per treatment for a total of 8 plots. In each block, a plot (ambient) experienced current ambient CO_2_ of ∼418 *μmol mol*^−1^, while the second plot was fumigated during daylight hours such that on average the target [CO_2_] within elevated plots was 600 *μmol mol*^−1^. This concentration represents the projected end of century concentration for the moderate shared socioeconomic pathway 4.5 (Arias *et al*., 2021). Monitoring of CO_2_ enrichment at the site showed that the [CO_2_] within elevated plots was within 10% of the target concentration 82% of the time during the growing season (Aspray *et al*., 2023).

#### PCR amplification and Sequencing of Leaf Samples

We collected leaf samples on 31 July 2020 from 15 haphazardly selected soybean plants per plot (total N = 120). The youngest fully expanded leaf for each plant was removed from each plant and kept on ice until returned to the lab. From each leaf we excised approximately 50mg tissue using a standard hole punch. Leaf discs were surface sterilized in 70% ethanol for 3 minutes followed by 0.8% bleach solution for 2 minutes and finally rinsed for 1 minute in sterile water. Sterilized leaf discs were flash frozen in liquid nitrogen and then stored at -80°C until DNA extraction.

Genomic DNA was extracted using the Qiagen Power Plant Pro DNeasy kit, following the manufacturer’s guidelines (Qiagen, Inc.). Fluidigm amplification followed by Illumina NovaSeq 6000 sequencing was performed at the W. M. Keck Center for Comparative and Functional Genomics at the University of Illinois, Urbana-Champaign. For fungi the internal transcribed spacer (ITS) region was amplified using primer pairs for ITS1 (ITS1F_ITS2, ITS1F 5’-CTTGGTCATTTAGAGGAAGTAA, ITS2 5’-GCTGCGTTCTTCATCGATGC) and ITS2 (ITS3_ITS4, 5’-GCATCGATGAAGAACGCAGC, ITS4 5’-TCCTCCGCTTATTGATATGC). For bacteria we used the 16S rRNA primer pair V4_515F_806R_New (V4_515F_New 5’-GTGYCAGCMGCCGCGGTAA, V4_806R_New 5’-GGACTACNVGGGTWTCTAAT).

We included two separate internal transcribed spacer (ITS) primers ITS1 and ITS2. In general, ITS2 amplifies a greater number of fungal taxa than ITS1 (e.g., Op De Beeck *et al*., 2014). Both have been used extensively as universal fungal primers yet these different sections amplify and differentiate separate taxa (Tekpınar and Kalmer, 2019).

#### Bioinformatics

We performed sample inference from amplicon data using the DADA2 denoising workflow (Callahan *et al*., 2016) in the statistical programming language R (R Core Team, 2022). We modified the standard ITS workflow (https://benjjneb.github.io/dada2/ITS_workflow.html) to only include forward reads as the quality of our reverse reads led to a higher proportion of data being discarded upon forward-reverse alignment. This is in line with suggestions that forward read-only approaches capture a more complete sample of the microbial community (Pauvert *et al*., 2019). Because the NovaSeq output bins error rates we used a modified error structure that enforced monotonicity and relaxed loess parameters which led to better error fitting curves.

DADA2 generates amplicon sequence variant (ASV) tables which allow for finer resolution than older OTU grouping methods (Callahan *et al*., 2017). We annotated ASVs using the UNITE sh_general_release_all_10.05.2021 database which included all eukaryotes for the ITS datasets (Abarenkov *et al*., 2023). We annotated bacterial sequence sets using the SILVA SILVA_SSU_r138_2019 database (Yilmaz *et al*., 2014). Sequences are deposited at GenBank (accession numbers xxxx-xxxx).

### Statistical analysis

#### Community analyses

Given that sequence read raw abundances are biased estimates of abundance and because microbial community data is compositional in nature, we calculated relative abundances of reads using Hellinger-transformed data (Legendre and Gallagher, 2001) before creating Bray-Curtis dissimilarity matrices used in further statistical procedures.

To test for differences in community composition between elevated and ambient [CO_2_] we used PERMANOVA with the ‘adonis2’ function in the ‘vegan’ package. Additionally, we visualized microbial communities in these treatments using non-metric multidimensional scaling (NMDS). We calculated Shannon diversity for elevated and ambient plots with the package ‘vegan’ (Oksanen *et al*., 2013).

#### Differential abundance analyses

To detect differences in abundance of reads across treatment groups, we estimated dispersion and logarithmic fold changes using the ‘DESeq2’ package. We tested for significant differences in log fold changes using Wald tests. We visualized these differences with the ‘EnhancedVolcano’ package as well as also plotting the log fold change shrinkage using the package ‘lfcShrink’ employing the apeglm method via the ‘apeglm’ package.

#### Network analyses

To understand how co-occurrence networks differed between the CO_2_ treatments we estimated co-occurrence networks using Sparse Inverse Covariance in the package ‘SpeiecEasi’ (Kurtz *et al*., 2023). To avoid spurious edges, we used the Meinshausen-Buhlmann estimation method. Before creating networks, we removed ASVs with low read counts (<10). This removed 25, 36, and 23 ASVs from the ITS1, ITS2, and 16s datasets, respectively.

We visualized the networks using the function ‘plot_network’ in the ‘phyloseq’ package (McMurdie and Holmes, 2013) visualizing both nodes and edges colored by their family for fungal datasets and their genus for the bacterial dataset. To test for statistical differences between elevated and control networks we used the package ‘NetCoMi’ (Peschel, 2023) which tests how networks differ in terms of network structure invariance, global strength invariance, and edge invariance using a permutational approach. To understand how roles within the networks differed between elevated and ambient CO_2_ conditions, we used ‘NetCoMi’ to report hub taxa where hub status is calculated by fitting a log-normal distribution to the centrality values to identify nodes with highest centrality values (Agler *et al*., 2016) though in this case we used centrality based on eigenvector scores. The similarity of taxa making up hub and most central nodes was statistically tested using the Jaccard index. All code and data for this work can be accessed on GitHub (https://github.com/connor-morozumi/soyleafendos.git).

## Results

### Overall

After trimming, removing chimeras, and bioinformatic denoising, we found a total of 835,777 reads corresponding to 360,468 reads within soybean leaves grown in ambient plots and 475,309 reads within elevated CO_2_ plots and for the ITS1 primer set. We found a total of 1,717,153 reads using the ITS2 primer set (752,887 in ambient plots and 964,266 in the elevated plots). For the bacterial dataset we found 260,679 total reads for the 16s primer set (113,846 reads for ambient plots and 146,833 reads for elevated plots). Classifying these sequences with existing taxonomic libraries, ambient CO_2_ plots had 872 ASVs whereas we found 1068 ASVs in elevated plots across all three primer sets.

### By taxa

In the ITS1 dataset we found 226 and 275 unique taxa across all plots in the control and elevated [CO_2_] treatments, respectively, with a per plot average of 93.8 ASVs in the control plots and 106 ASVs in the elevated plots. In the ITS2 dataset we found 533 and 664 unique taxa across all plots in the control and elevated [CO_2_] treatments, respectively, with a per plot average of 175.8 ASVs within the control plots and 214 ASVs within the elevated plots. The ITS1 and ITS2 datasets differed from one another. The ITS1 dataset contained 31 genera found not present in the ITS2 dataset while, similarly, the ITS2 dataset contained 33 unique genera not found in the ITS1 dataset. At the family level, 15 families not detected by the ITS2 primer set were found in the ITS1 dataset. The ITS2 primer set found 20 unique families absent in the ITS1 dataset. In sum, at the genus level, 47.81% taxa were found in both primers while at the family level this equated to 55.70% of taxa being shared (Supplementary Figure S1, Supplementary Figure S2). We present visualizations for community composition and networks for ITS2 and 16S here while analogous results for ITS1 can be found in the Supplementary Materials (Supplementary Figure S3, Supplementary Figure S4).

Fungal community composition differed across CO_2_ treatment for both ITS1 (PERMANOVA , R^2^= 0.015, p = 0.034, S3) and ITS2 (PERMANOVA, R^2^= 0.031, p = 0.001, Figure 1A). For ITS1, Shannon diversity was similar across treatments, S= 1.81 and S= 1.81 for ambient and elevated treatments, respectively. Shannon diversity was unchanged across treatments for the ITS2 primer as well (S= 2.16, S= 2.18 for ambient and elevated [CO_2_], respectively). For the ITS2 primer set, we found statistically significant differential abundance changes for 139 ASVs which represents 13.31% of the total ASVs present (log fold change adjusted for false discovery rate, p-value < 0.1, Supplementary Figure S5, Supplementary Table S1). Analogous differential abundance results for the ITS1 primer can be found in Supplementary Data (Supplementary Figure S6, Supplementary Table S2).

**Figure 1.**
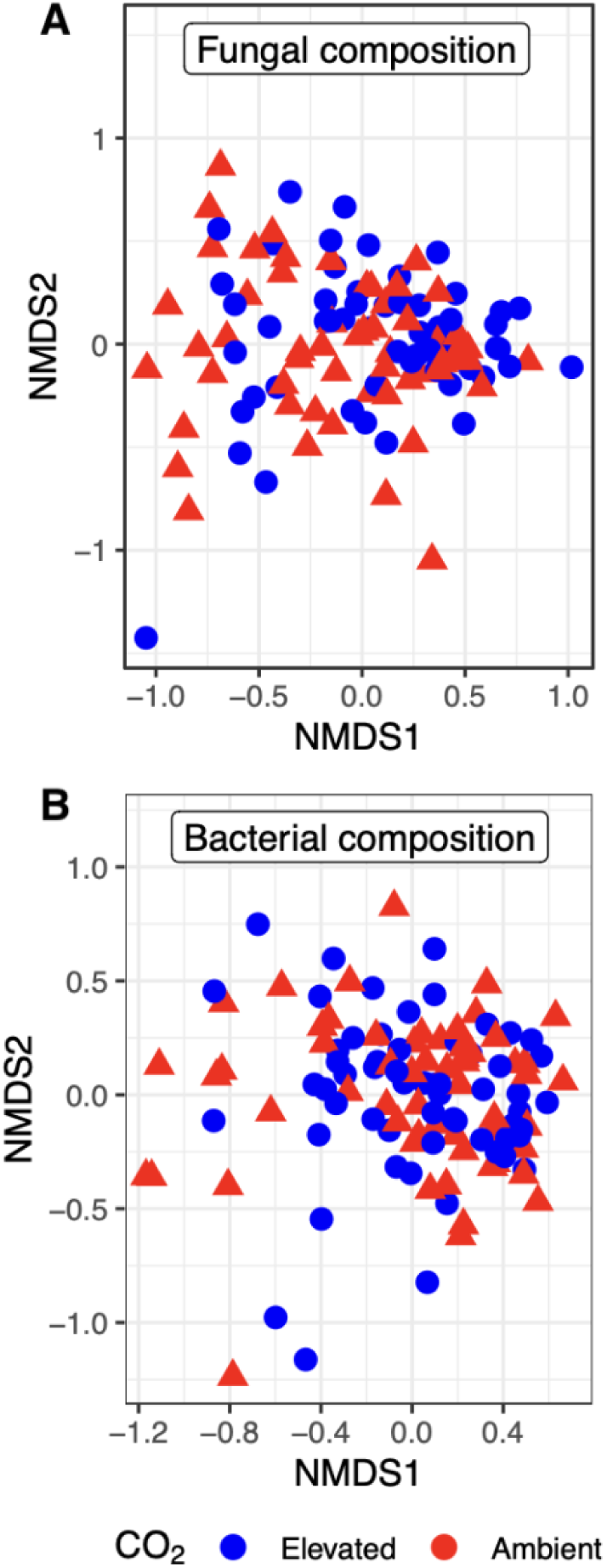
**A.** Soybean foliar fungal composition differed between elevated (red triangles) and ambient (blue circles) CO_2_ treatments (PERMANOVA, R^2= 0.031, p =0.001). Shown are community composition along NMDS1 and NMDS2 based on ITS2 primer set, community composition results were largely analogous for the ITS1 primer set. **B.** Soybean foliar bacterial composition based on the 16S primer set did not differ between elevated and ambient [CO_2_] treatments (PERMANOVA, R^2= 0.012, p =0.1). Alt text: Two panels sit atop one another. They both are graphed the same. The axes for both are labeled NMDS 1 and NMDS 2 for the x and y axes, respectfully. In the plot are points in solid fill. There are blue circles and red triangles each representing the microbial composition of each plot. For the top graph (fungal composition), the points are centered at 0,0. The spread of all points ranges from about -1 to +1 on each axis with slightly more spread on axis 1 than axis 2. While there is a large overlap in the points there is more red triangles to the left side of the x axis (NMDS 1) and a greater concentration of blue circles to the right side of NMDS 1. There is a blue outlier high on NMDS 2 in the upper lefthand corner. Other than this outlier there is a tighter dispersion of points for the blue circles than the red triangles which are spread across the graph. For the bottom graph (bacterial composition), the points are centered at -0.5,0. The spread of most points ranges from about -1 to +1 on axis 1 and from -1 to +0.5 on axis 2. While there is a large overlap in the points there is more red triangles to the left side of the x axis (NMDS 1) and a greater concentration of blue circles to the right side of NMDS 1. There are more outliers in this graph than in the top panel: two blue circles and one red triangle have low NMDS 2 values less than -1. The dispersion of points for the blue circles and the red triangles are similarly spread across the graph.

Using the 16s primer set, we found 113 and 129 unique bacterial taxa across all plots in the control and elevated [CO_2_] treatments, respectively. Per plot, we found an average richness of 46.25 ASVs in the control plots and 51.50 ASVs in the elevated plots. Bacterial composition did not differ across CO_2_ treatments (PERMANOVA, R^2^= 0.012, p = 0.16, Figure 1B). Shannon diversity was similar across the treatments (S=1.14, S= 1.19 for ambient and elevated [CO_2_], respectively). For the 16S primer set, 14 ASVs had statistically significant changes in differential abundances (6.54% of the total ASVs present (log fold change adjusted for false discovery rate, p-value < 0.1, Supplementary Figure S7, Table S3).

### Networks

Fungal networks differed between the CO_2_ treatments. Ambient and elevated fungal networks based on the ITS2 primer pair had a similar proportion of total links (ambient edge density = 0.041; elevated edge density = 0.034) and negative links (ambient negative links = 1.66%; elevated negative links = 1.51%;) (Figure 2, Table 1). At the node level, network structural metrics of centrality indicated turnover such that dominant and highly connected taxa differed between the treatments (taxa with highest eigenvector centrality: J= 0.02, p<0.001, hub taxa: J = 0.00, p < 0.001). Hub taxa identified in the ITS2 dataset differed between the CO_2_ treatment groups. Seven *Colletotrichum* strains (6 of *Colletotrichum spaethianum* and 1 of *Colletotrichum cliviae* were identified to be hubs in ambient [CO_2_] but not in elevated [CO_2_] (no *Colletotrichum* strains were hubs in elevated plots.) Across the treatment groups several *Cladosporium* strains were identified as hubs (Table 2). Networks visualized for the ITS1 primer set can be found in Supplementary Figure S2. We plotted community composition with influential hub species identified which can be found in Supplementary Figure S8.

**Figure 2.**
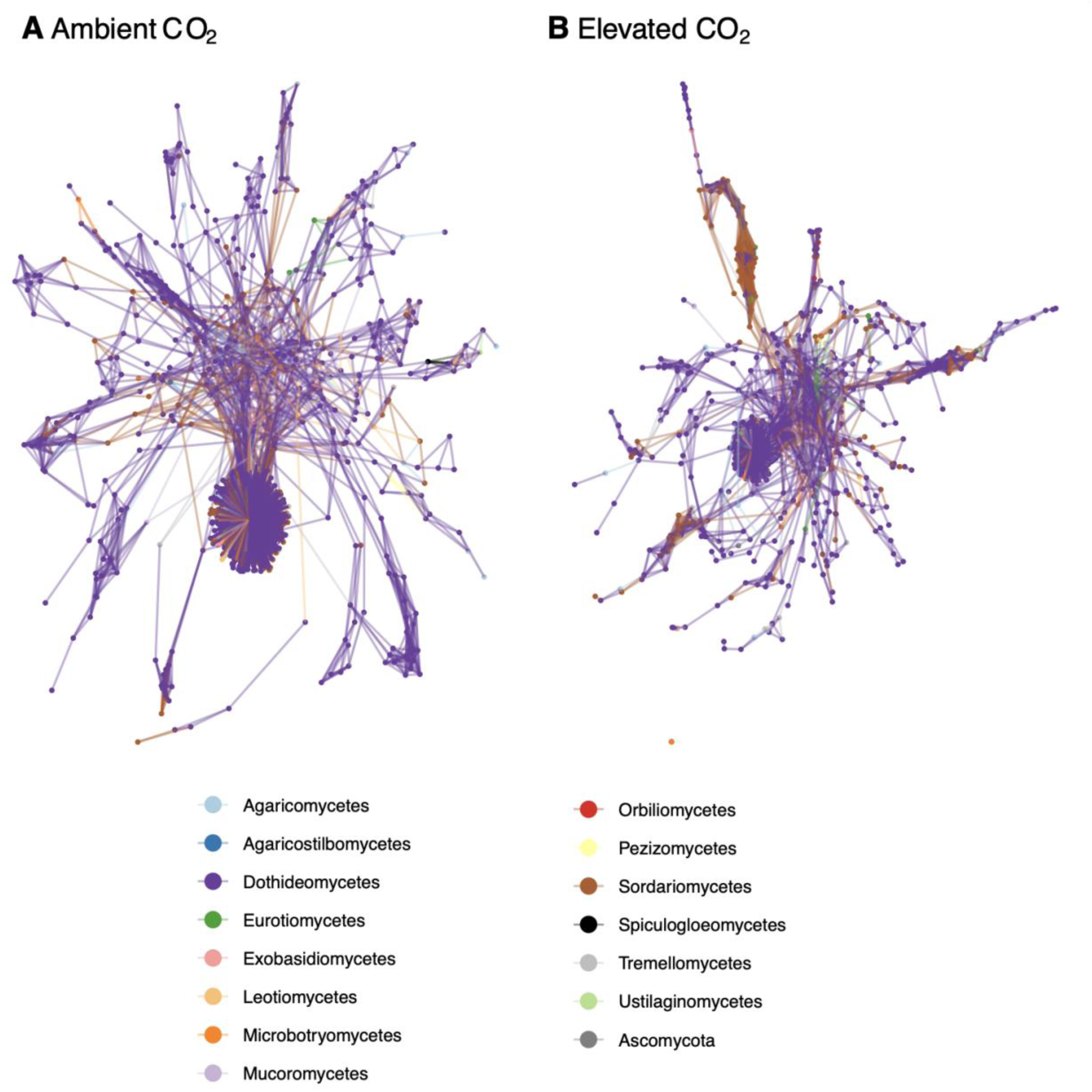
Fungal co-occurrence networks differed between foliar taxa from soybean plants grown in ambient(A) and elevated(B) [CO_2_] conditions identified with the ITS2 primer set. Nodes and edges colored by fungal order. A two-panel figure with the panels side by side. Each show a colorful network of points (nodes) and lines (links) connecting the nodes. The nodes are colored by the order for each taxon in the network. The left (A) panel is labeled Ambient CO_2_ while the right (B) panel is labeled Elevated CO_2_. Each network has a densely connected portion that looks like a ball with spikes on it. This densely connected portion is linked more diffusely with a second less dense region that still has many nodes graphed close together which are then connected to the rest of the network that radiates out away from the more central taxa. In both panels all nodes are connected to one another such that there are no smaller isolated networks. In the left Ambient panel there are more links from the main center ball to the second region while in the right panel (elevated CO_2_) there are less links connecting these central taxa to the rest of the network. Below both panels there is a legend listing all the family names.

**Table 1.** Network Metrics.

| Metric | Ambient | Elevated | Difference |
| --- | --- | --- | --- |
| **ITS2** |  |  |  |
| Number of components | 1.000 | 2.000 | 1.000 |
| Clustering coefficient | 0.706 | 0.641 | 0.071 |
| Modularity | 0.149 | 0.330 | 0.174 |
| Positive edge percentage | 98.345 | 98.488 | 0.072 |
| Edge density | 0.041 | 0.034 | 0.006 |
| **16S** |  |  |  |
| Number of components | 18.000 | 11.000 | 7.000 |
| Clustering coefficient | 0.683 | 0.587 | 0.095 |
| Modularity | 0.255 | 0.345 | 0.090 |
| Positive edge percentage | 96.330 | 90.957 | 5.374 |
| Edge density | 0.030 | 0.032 | 0.002 |
| <sup>1</sup> Number of components = total number of separate sets of connected nodes<br><sup>2</sup> Clustering coefficient= measures the local group cohesiveness and is calculated as the average of the fraction of connected neighbors for every vertex<br><sup>3</sup> Modularity = the degree to which nodes are separated into distinct groups. Higher modularity scores indicate dense connections between the nodes within modules but sparse connections between nodes in different modules |  |  |  |

**Table 2.** Hub Taxa by Primer Set.

| Ambient | Elevated |
| --- | --- |
| **16s** |  |
| *Hymenobacter* sp. 1 | *Aurantimonas* sp. 2 |
| *Methylobacterium-Methylorubrum* sp.<br>7, 10, 11, 12, 14, 17, 35 | *Aureimonas* sp.<br>2, 7 |
| *Sphingomonas mucosissima* 1 | *Escherichia-Shigella* sp. 2 |
|  | *Methylobacterium-Methylorubrum* sp.<br>5, 6, 9 |
|  | *Oscillatoria* sp. 26 |
|  | *Quadrisphaera* sp. 1 |
|  | *Sphingomonas* sp. 2 |
| Total hubs identified= 9 | Total hubs identified= 10 |
| **ITS2** |  |
| *Alternariaster* sp. 1, 2, 3 | *Alternaria alternata*<br>2, 3 |
| *Cladosporium dominicanum*<br>1, 2 | *Cladosporium grevilleae*<br>2, 6 |
| *Cladosporium grevilleae* 3 | *Cladosporium ramotenellum*<br>3, 6, 15, 18, 21, 59, 66 |
| *Cladosporium ramotenellum*<br>3, 4, 5, 7, 9, 10, 12, 61 | *Coniothyrium* sp.<br>2, 3 |
| *Cladosporium* sp. 1 | *Diaporthe* sp. 1 |
| *Colletotrichum cliviae* 2 | *Didymella* sp.<br>4, 9, 10, 30 |
| *Colletotrichum spaethianum*<br>2, 3, 4, 5, 6, 7, 24, 25 | *Didymellaceae*<br>6, 10 |
| *Diatrypaceae* 2 | *Didymosphaeriaceae*<br>1, 18 |
| *Didymella* sp.<br>2, 5, 6, 7 | *Dissoconium* sp. 1 |
| *Didymellaceae* 8, 37 | *Epicoccum* sp.<br>3, 7, 10, 12, 16 |
| *Didymocyrtis cladoniicola* 3 | *Fusarium* sp. 4 |
| *Epicoccum* sp.<br>2, 4, 5 | *Gibberella intricans*<br>2 |
| *Epicoccum thailandicum*<br>1 | *Gibellulopsis chrysanthemi* 1 |
| *Fusarium* sp. 3 | *Hannaella coprosmae* 1 |
| *Hannaella oryzae*<br>2, 3 | *Helotiales* 1 |
| *Hannaella zeae* 1 | *Keissleriella quadrisepata* 2 |
| *Keissleriella quadrisepata* 3 | *Keissleriella* sp. 1 |
| *Microdochium colombiense* | *Keissleriella taminensis* 2 |
| *Neosascochyta argentina* 2 | *Leptospora* sp. 5 |
| *Neosetophoma* sp. 9 | *Murispora* sp.<br>2, 3 |
| *Orbiliales*<br>1, 2 | *Neosascochyta europaea* 1 |
| *Paraconiothyrium* sp. 3 | *Neodevriesiaceae* 1 |
| *Penicillium bialowiezense* 1 | *Neosetophoma* sp. 1<br>8, 11, 27 |
| *Pleosporales* 10 | *Nigrospora* sp. 1 |
| *Preussia tenerifae* 1 | *Periconia* sp. 1 |
| *Septoria* sp. 2 | *Phaeosphaeriaceae*<br>1, 2 |
| *Talaromyces trachyspermus* 1 | *Setophaeosphaeria badalingensis* 1 |
| *Tetraplospheariaceae* 1 | *Stagonosporopsis* sp. 1 |
|  | *Stagonosporopsis valerianellae* 1 |
|  | *Xylodon erastii* 1 |
| Total hubs identified= 50 | Total hubs identified=50 |

Bacterial networks also differed between the elevated and ambient [CO_2_] (Figure 3, Table 1). Bacterial networks in the elevated treatment had similar total links (elevated edge density = 0.041; ambient edge density = 0.050), and greater proportion of negative links (elevated = 9.34%; ambient = 3.90%). Additionally, we observed similar turnover of highly connected taxa across the treatment groups (taxa with highest eigenvector centrality: J= 0.00, p< 0.001; hub taxa: J = 0.00, p < 0.001). In elevated plots one ASV of *Aurantimonas sp.* and two ASVs of *Aureimonas* sp. were identified as hub taxa, as were three ASVs of *Methylobacterium* sp. and one ASV of *Oscillatoria* sp. In the ambient plots, hub taxa were dominated by seven different ASVs identified as *Methylobacterium* sp. along with two taxa not identified as hubs in the elevated plots: *Sphingomonas mucosissima* and *Hymenobacter* sp.

**Figure 3:**
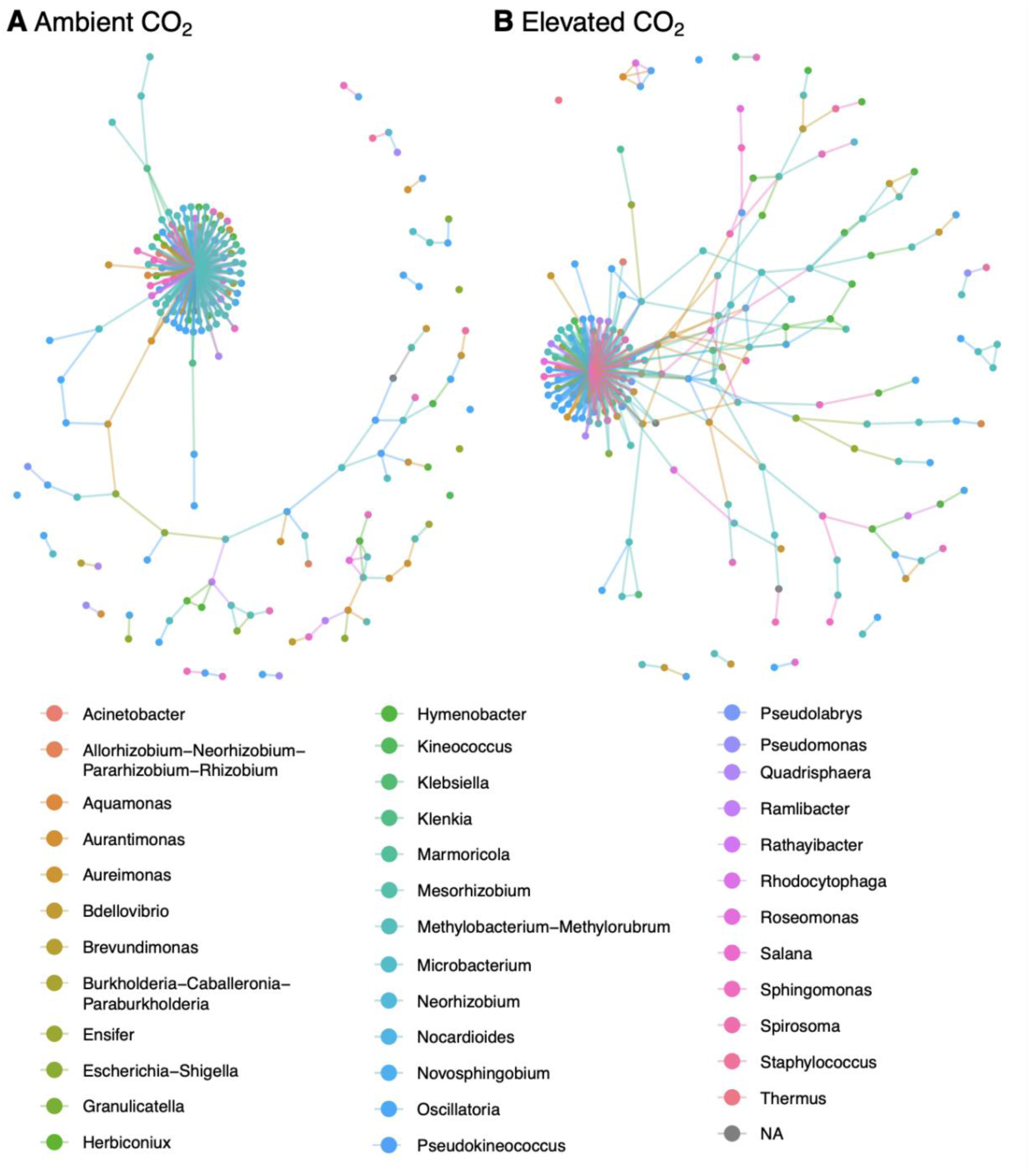
Networks show different co-occurrence patterns for foliar bacterial taxa from soybean plants grown in ambient(A) and elevated(B) [CO_2_] conditions identified with the 16s primer set. Nodes and edges colored by bacterial genus. A two-panel figure with the panels side by side. Each show a colorful network of points (nodes) and lines (links) connecting the nodes. The nodes are colored by the genera for each taxon in the network. The left (A) panel is labeled Ambient CO_2_ while the right (B) panel is labeled Elevated CO_2_. Each network has a densely connected portion that looks like a ball with spikes on it. This densely connected portion is linked diffusely with the rest of the network that radiates out away from the ball. There are more nodes that are connected only to their small network and not the main part of the network in the left panel. In the left Ambient panel, there are only a few connections or links to the main center ball while in the right panel (elevated CO_2_) there are more links connecting these central taxa to the rest of the network. Below both panels there is a legend listing all the genera names.

## Discussion

In this study we compared leaf endophytic fungal and bacterial communities in soybean grown under elevated and ambient [CO_2_], thus testing how these microbiomes might shift under a future climate change scenario. Using a metabarcoding approach, we found that fungal community composition in soybean leaves was sensitive to [CO_2_] while bacterial communities were more stable across the experimental treatments. Additionally, we found marked differences in co-occurrence networks for both fungal and bacterial taxa especially in terms of which species constituted the hub within each treatment. We found no strong directional changes towards more negative interactions as we had predicted for fungal networks while bacterial networks had slightly more negative interactions under elevated [CO_2_]. Contrary to our predictions, networks under elevated [CO_2_] were also similar in the number of links as their ambient controls. Taken together our findings indicate that endophyte communities within soybean will change under future climate change.

We expected certain taxa to be able to take advantage of elevated [CO_2_] and potentially outcompete other microbes in soybean leaves. This is because elevated CO_2_ is associated with increased leaf carbohydrates (Ainsworth *et al*., 2002; Gonçalves *et al*., 2021) that we thought would be capitalized on by select dominant taxa . Thus, we had hypothesized that co-occurrence networks within elevated CO_2_ conditions would have fewer total links and a higher proportion of negative co-occurrence correlations among the remaining taxa. Instead, we found that fungal networks under elevated [CO_2_] had similar number of total links and that the proportion of positive links was unchanged, contrary to our hypotheses. The stress gradient hypothesis predicts that more stressful conditions act to strengthen positive interactions (He *et al*., 2013); therefore, it may be that increased CO_2_ represents a stressor, which countered the effects of dominant competitors. For bacterial networks, we did not find differences in the density of links between the two CO_2_ treatments though we did find a higher proportion of negative links under elevated CO_2_ conditions. While co-occurrence of taxa cannot be interpreted directly as a proxy for species interaction type, our finding that networks were not becoming dominated by negative links may indicate that associations between taxa are not becoming more antagonistic under elevated [CO_2_]. More highly connected communities are associated with higher stability (Herren and McMahon, 2017) so finding that elevated [CO_2_] is not perturbing our networks in terms of link densities may mean microbial communities in soybean leaves are resistant to changes due to increased [CO_2_], though ecological interpretation of network properties such as stability is still poorly understood.

We found bacterial communities to be largely stable under elevated [CO_2_], which was consistent with previous studies. For example, bacterial communities in rice leaves were only sensitive to elevated [CO_2_] in combination with additional experimental manipulation of temperature, rather than elevated [CO_2_] alone (Ren *et al*., 2015). Similarly, in a culture-based study of cotton using a FACE experiment, bacterial communities were unchanged under elevated [CO_2_] (Runion *et al*., 1994). The bacterial community we found in soybean leaves in both elevated and ambient plots was consistent with previous studies of other species. For example, across both of our treatment groups we found ca. 65% of our bacterial ASVs were in class Alphaproteobacteria (phylum Proteobacteria) which is a major player in microbiomes of many plant species (Müller *et al*., 2016). Research on foliar bacterial change in response to CO_2_ manipulation is very scant and the current work appears to be some of the only to test such changes in soybean.

Previous work from SoyFACE found that elevated [CO_2_] decreased the abundance of a strain of *Methylobacterium* sp., and that this strain acted as an antagonist to fungal endophytes *in vitro* (Christian *et al*., 2021). We found the same *Methylobacterium* strain when we assessed soybean microbiomes with culture-independent sequencing methods as well as about 50 additional *Methylobacterium* ASVs. We found differences in abundance of two *Methylobacterium* strains between elevated and ambient [CO_2_] and multiple *Methylobacterium* strains played more central roles in bacterial networks within ambient plots compared to elevated plots. Central taxa within networks are thought to help shape the remaining microbial community (Agler *et al*., 2016).

Given that these *Methylobacterium* mainly have antagonistic relationships with other fungal endophytes in soybean (Christian *et al*., 2021) and reduce pathogenic effects in other systems (Madhaiyan *et al*., 2006; Dourado *et al*., 2015) their loss as hub taxa could have notable downstream effects under elevated [CO_2_] if they are playing disease suppression roles in soybean. Indeed Sosa Marquez *et al*. (2025) found that this strain is highly beneficial for soybean, at least under axenic conditions in the greenhouse.

The relative stability in community composition we found in bacterial microbiomes in the leaves in response to elevated [CO_2_] differed from what has been found in root microbiomes. Within the same SoyFACE experimental array as the current study, bacterial communities in the rhizosphere of soybean differed across CO_2_ treatments, though the communities found within roots were unaffected (Wang *et al*., 2017). This study also found a decrease in bacterial diversity in the rhizosphere associated with elevated [CO_2_]. Similarly, bacterial communities of the rhizosphere of soybean differed across CO_2_ treatments (Yu *et al*., 2016). Conversely, other studies have found bacterial communities in roots to be stable in elevated [CO_2_]. For example, one study in ryegrass found that soil type structured bacterial communities in roots to a far greater degree than CO_2_ fumigation (Chen *et al*., 2016). Conversely, other studies have found bacterial communities in roots to be stable in the face of elevated [CO_2_] (Jossi *et al*., 2006; Lesaulnier *et al*., 2008). We found modest bacterial diversity in leaves—in line with other studies of the phyllosphere of soy (e.g., Chen *et al*., 2022). It may be that the abiotic conditions necessary for surviving aboveground in leaves drives bacterial diversity and composition to a greater extent than [CO_2_].

We found different fungal communities within soybean leaves grown in elevated versus ambient [CO_2_]. This mirrors previous research from our group which found culturable fungal leaf endophyte communities differed between elevated and ambient plots at SoyFACE (Christian *et al*., 2021). Our results also are in line with previous research which found that fungal communities in the rhizosphere of soybean are similarly responsive to elevated [CO_2_] (Yu *et al*., 2018). While documenting shifts in the composition of the microbiome is important, it is also critical to assess how microbial interactions within these systems are being altered.

Our networks showed that elevated [CO_2_] acted to shift co-occurrence patterns within soybean leaves. Our findings contrast past work which found simplified fungal networks under elevated [CO_2_] in the rhizosphere (Yu *et al*., 2018) though this was not the case for either fungal or bacterial networks of endospheric and rhizospheric taxa found in wheat and rice grown in a field-based study also using FACE technology (Gao *et al*., 2022). In the current study, taxa playing central roles within our fungal networks were different between the [CO_2_] treatments. Within ambient plots, *Colletotrichum* ASVs were important to network structure whereas they were not identified as hubs in elevated plots. This may represent an important turnover within elevated plots given 97% of ASVs that we classified as *Colletotrichum spaethianum* were >99% similar to a strain found to be competitive with key *Methylobacterium* sp. (Christian *et al*., 2021).

Intriguingly, whereas the previous study found elevated [CO_2_] increased the abundance of *Colletotrichum spaethianum*, in our study this strain is playing more a central role in the network under ambient conditions. In both elevated and ambient [CO_2_] in the current study, several yeast taxa played hub roles. Little is known about the functional roles of yeasts in plant microbiomes (Gouka *et al*., 2022). One hub species in elevated [CO_2_] conditions was the likely fungal pathogen *Coniothyrium* sp. which was not central to networks within ambient conditions.

Bacterial hub taxa were also different across the [CO_2_] treatment. Enrichment of *Aurantimonas* sp. and closely related *Aureimonas* spp. in the suite of taxa playing central roles in the elevated bacterial networks is particularly interesting as these are phylogenetically close to rhizobia. Both *Aurantimonas* and *Methylobacterium* found within aboveground stems have been found to be sensitive to N fertilization and nodulation phenotype in soybean (Ikeda et al. 2010) (Ikeda *et al*., 2010) and it is notable that we found them to be responding to [CO_2_] differentially within leaves.

The entire ITS region is too long for Illumina-based sequencing platforms, so researchers commonly sequence either the ITS1 or ITS2 region. However, there is still a lack of consensus among researchers on the optimal fungal metabarcode, and a continued need for studies that directly compare primer sets for both the ITS1 and ITS2 regions (Winand *et al*., 2025). We found far fewer taxa using the ITS1 primer set compared to the ITS2 gene region. Tests of these primer sets have indicated that these two primers introduce different biases. The ITS1 primer set we used is known to suffer from a high degree of mismatches which may necessitate relaxing the PCR conditions in order to amplify all fungi in a sample (Bellemain *et al*., 2010). This same *in silico* study also found that targeting the ITS2 region preferentially amplifies ascomycetes while targeting the ITS1 region is biased toward basidiomycetes (Bellemain *et al*., 2010).

There are a few caveats and limitations to the work we present here. We collected plant tissues only once during the growing season. Thus, this represents a snapshot of a community known to have considerable seasonal turnover (Copeland *et al*., 2015) which may affect community composition and network differences across our treatments. Additionally, we only tested the effects of one type of global climate change—the increased concentration of atmospheric CO_2_. Experimentally crossing this discrete stressor with others associated with climate change (e.g., increased ambient temperatures) is needed in our system. It may be that the impacts of elevated [CO_2_] do not fully manifest until paired with other climate change stress. For example, Ren et al. (2015) found that the effects of elevated [CO_2_] did not begin to change rice microbiomes until it was manipulated in combination with temperature. These kinds of tests will be especially important given the multiple simultaneous stressors that plants and their microbial partners face in a rapidly changing world. Another caveat to note is that network co-occurrences are not necessarily indicative of species interactions. Careful *in vitro* and *in vivo* work stemming from network-based predictions, such as the tests of disease mitigation by the suite of *Methylobacterium* strains (e.g., Christian *et al*., 2021), will be important steps towards a more functional understanding of the leaf microbiome. Our work provides one of the first investigations into how global climate change may induce alterations to leaf microbiomes and thus some of these limitations might better be thought of as fruitful next directions in furthering our understanding of how microbial communities within crop systems will fare under anthropogenic change.

## Conclusion

Responses of ecosystems to anthropogenic change are likely to be dependent, in part, on microbial responses to these stressors. Microbes that make up plant microbiomes, in particular leaf endophytes, are known to play protective roles in their host especially in response to environmental stressors. We found that communities of endophytes in leaves—especially fungal taxa—were sensitive to elevated [CO_2_] likely to be reached under moderate scenarios of ongoing climate change. Unanswered questions remain and future work should investigate how taxa most sensitive to these perturbations play a role in host function.

## Supplementary data

The following supplementary data are available online.

Figure S1: Taxa overlap at the family level between the datasets generated with the ITS1 vs ITS2 primer sets.

Figure S2: Taxa overlap at the genus level between the datasets generated with the ITS1 vs ITS2 primer sets.

Figure S3. Soybean foliar fungal composition based on the ITS1 primer set.

Figure S4. Fungal co-occurrence networks based on the ITS1 primer set.

Figure S5. Differential abundance for fungal ASVs detected with the ITS2 primer set.

Table S1. Fungal taxa identified with the ITS2 primer set with statistically significant log fold change.

Figure S6. Differential abundance shows log fold change for fungal ASVs detected with the ITS1 primer set.

Table S2. Fungal taxa identified with the ITS1 primer set with statistically significant log fold change.

Figure S7. Differential abundance shows log fold change for bacterial ASVs detected with the 16S primer set.

Table S3. Bacterial taxa identified with the 16s primer set with statistically significant log fold change.

Figure S8. Community composition of fungal and bacterial ASVs with hub taxa that had influential NMDS scores labeled.

## Acknowledgements

This work would not have been possible without the field and lab assistance of Karla Griesbaum, Ivan Sosa Marquez, and Essie Steinbacher who assisted with sample collection and processing.

Additionally, the Keck Center at UIUC performed the sequencing. We are indebted to all researchers and crew at SoyFACE for establishing and maintaining the experimental array.

## Data Availability

Sequence data underlying this article are available in GenBank at [*updated url upon submission*] and can be accessed with accession numbers xxxx-xxxx. All other data to support the findings of this study are openly available in GitHub at [https://github.com/connor-morozumi/soyleafendos.git].

