## Supplementary figures for "Elevated CO_2_ induces phyllosphere community changes in soybean"

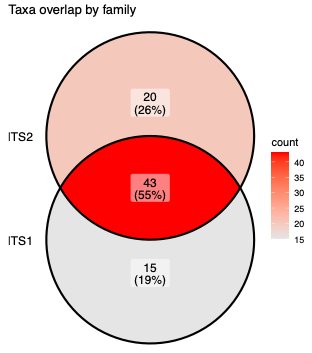
Figure S1: Taxa overlap at the family level between the datasets generated with the ITS1 vs ITS2 primer sets.

Figure S2: Taxa overlap at the genus level between the datasets generated with the ITS1 vs ITS2 primer sets.

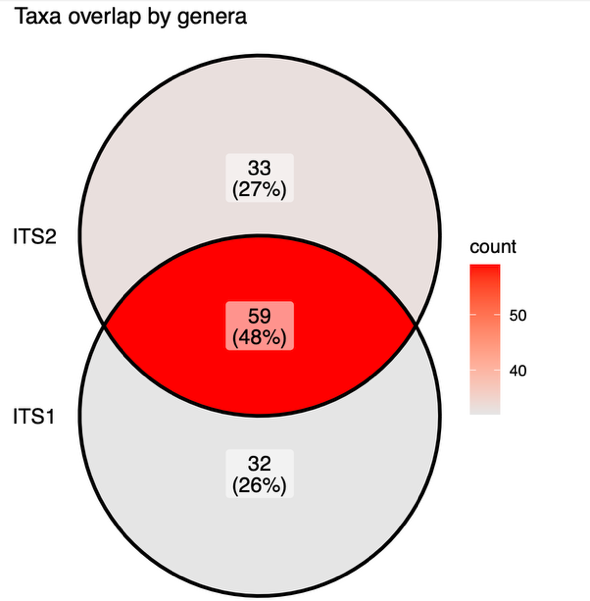

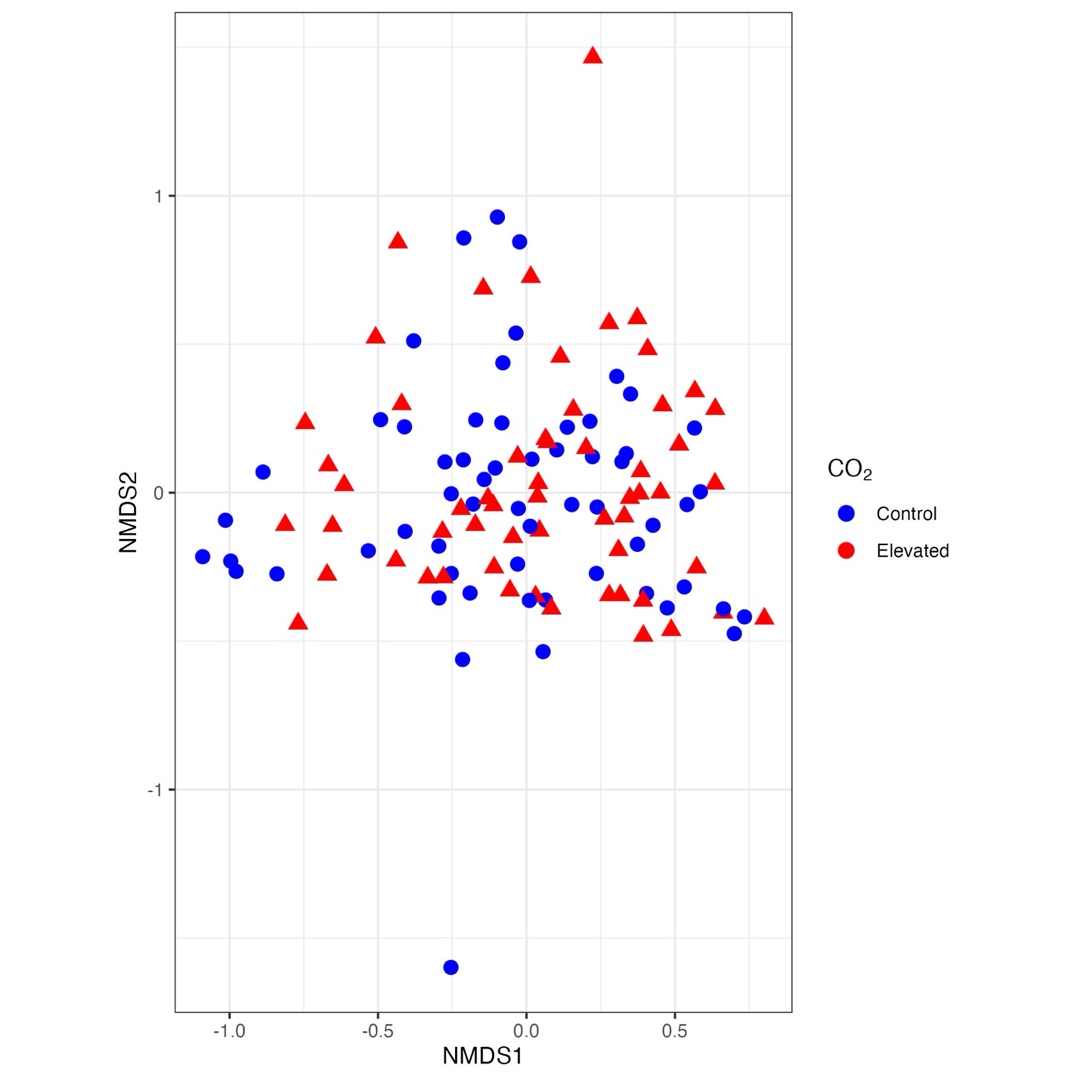

Figure S3. Soybean foliar fungal composition differed between elevated (red triangles) and ambient (blue circles) CO_2_ treatments (PERMANOVA , R^2^= 0.015, p = 0.034). Shown is community composition along NMDS1 and NMDS2 based on the ITS1 primer set.

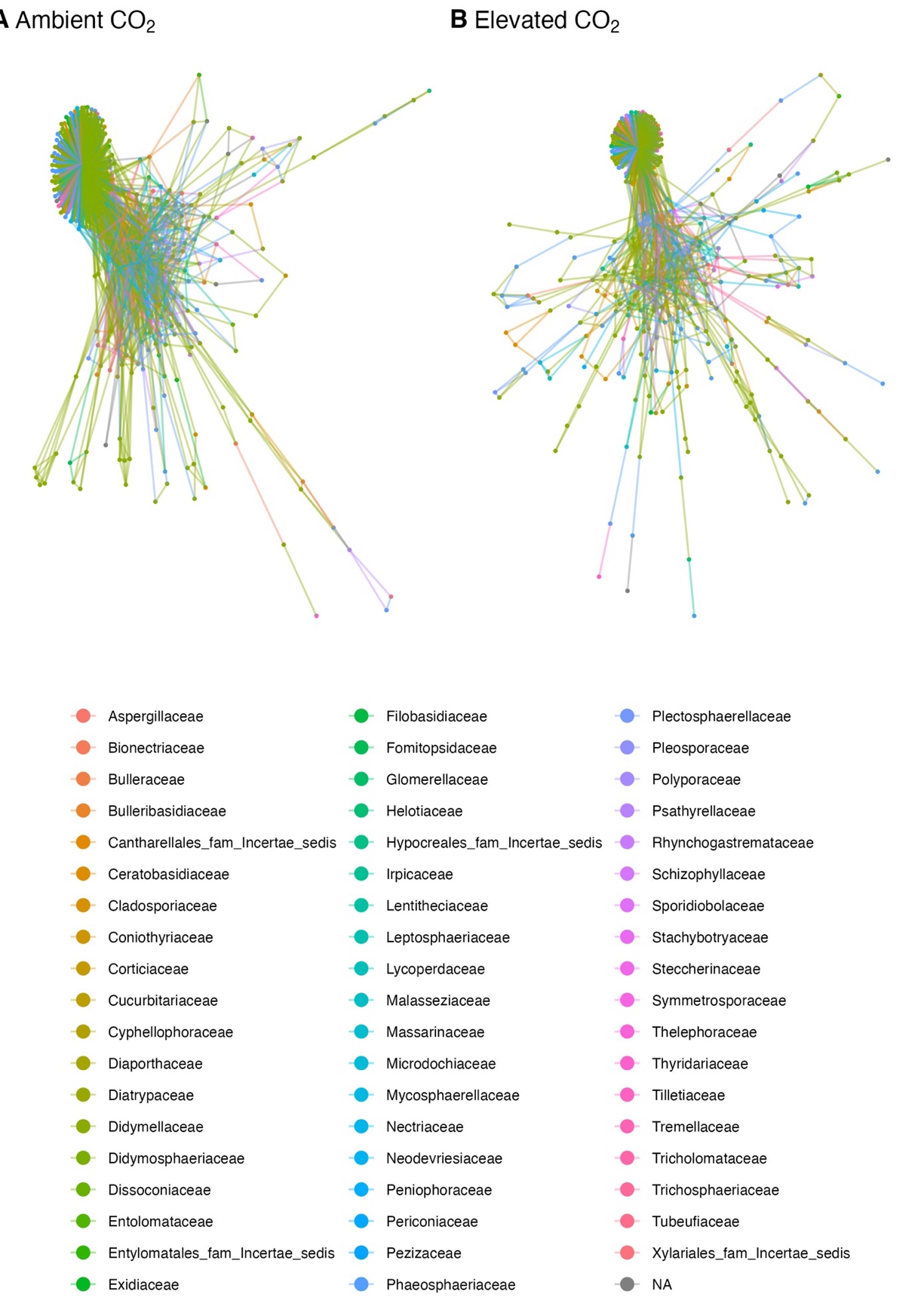

Figure S4. Fungal co-occurrence networks differed between foliar taxa from soybean plants grown under ambient(A) and elevated(B) [CO_2_] conditions. Fungal DNA was amplified using the ITS1 primer set. Nodes and edges colored by fungal family.

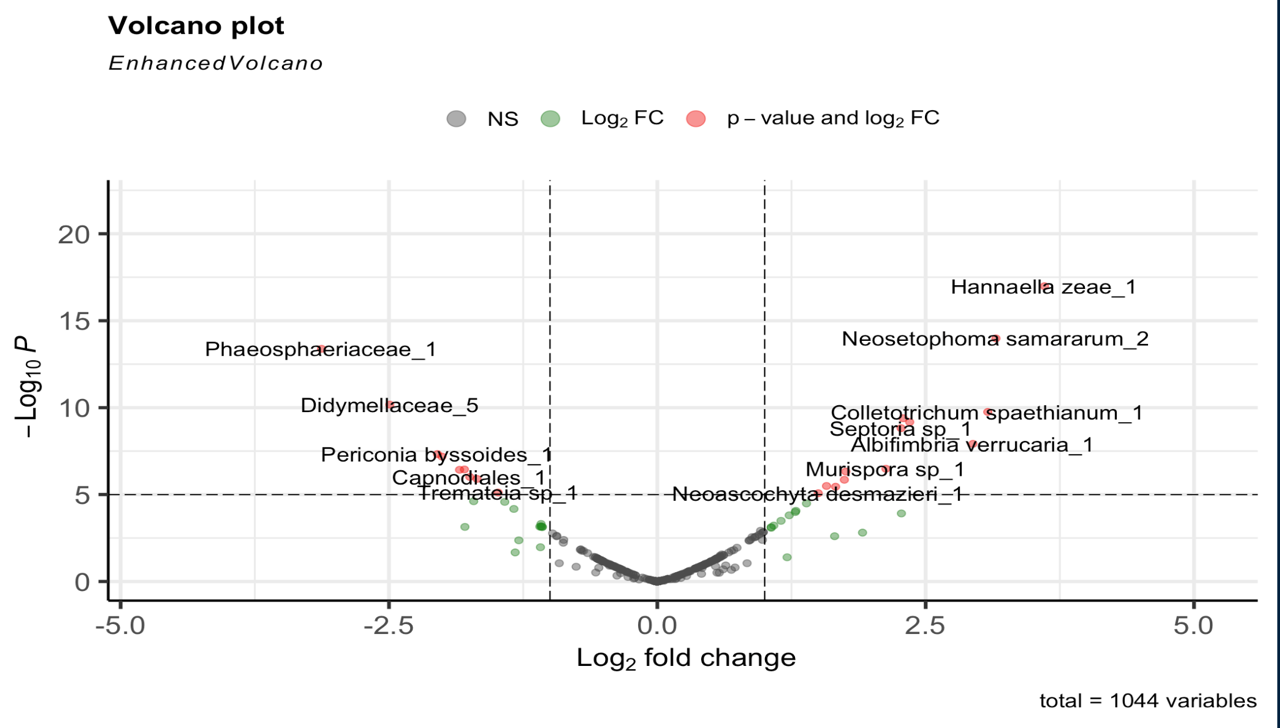

Figure S5. Differential abundance shows log fold change for fungal ASVs detected with the ITS2 primer set. ASVs with positive values to the right of 0 on the x-axis are found in greater abundance in the elevated CO_2_ plots whereas ASVs to the left are found in greater abundance in the control (ambient CO_2_) plots.

Table S1. Fungal taxa identified with the ITS2 primer set with statistically significant log fold change. List has been further subset to those ASVs with log fold change scores >1. Positive log fold change scores indicate higher abundance in the elevated CO_2_ plots whereas negative scores indicate higher abundance in the control/ambient CO_2_ plots.

|  | Base Mean | log2  Fold Change | lfcSE | stat | pvalue | padj |
| --- | --- | --- | --- | --- | --- | --- |
| Hannaella zeae_1 | 6.803 | 3.607 | 0.421 | 8.570 | 0.000 | 0.000 |
| Neosetophoma samararum_2 | 5.026 | 3.154 | 0.408 | 7.735 | 0.000 | 0.000 |
| Phaeosphaeriaceae_1 | 5.099 | -3.129 | 0.414 | -7.556 | 0.000 | 0.000 |
| Didymellaceae_5 | 3.465 | -2.492 | 0.382 | -6.527 | 0.000 | 0.000 |
| Colletotrichum spaethianum_1 | 15.871 | 3.078 | 0.483 | 6.380 | 0.000 | 0.000 |
| Epicoccum thailandicum_1 | 3.046 | 2.291 | 0.366 | 6.251 | 0.000 | 0.000 |
| Pleosporales_1 | 3.100 | 2.351 | 0.381 | 6.172 | 0.000 | 0.000 |
| Septoria sp_1 | 3.011 | 2.271 | 0.376 | 6.043 | 0.000 | 0.000 |
| Albifimbria verrucaria_1 | 29.231 | 2.940 | 0.516 | 5.698 | 0.000 | 0.000 |
| Periconia byssoides_1 | 2.683 | -2.048 | 0.375 | -5.462 | 0.000 | 0.000 |
| Neosetophoma sp_4 | 2.576 | -2.007 | 0.371 | -5.404 | 0.000 | 0.000 |
| Murispora sp_1 | 5.229 | 2.130 | 0.417 | 5.107 | 0.000 | 0.000 |
| Neoascochyta sp_1 | 2.337 | -1.797 | 0.353 | -5.089 | 0.000 | 0.000 |
| Ophiosphaerella aquatica_2 | 2.807 | -1.843 | 0.363 | -5.080 | 0.000 | 0.000 |
| Plectosphaerella cucumerina_1 | 2.265 | 1.756 | 0.349 | 5.036 | 0.000 | 0.000 |
| Capnodiales_1 | 2.274 | -1.747 | 0.357 | -4.898 | 0.000 | 0.000 |
| Epicoccum sp_11 | 2.194 | -1.680 | 0.347 | -4.847 | 0.000 | 0.000 |
| Blakeslea trispora_3 | 3.010 | 1.742 | 0.361 | 4.823 | 0.000 | 0.000 |
| Cladosporium ramotenellum_20 | 2.061 | 1.576 | 0.339 | 4.656 | 0.000 | 0.000 |
| Blakeslea trispora_2 | 3.029 | 1.661 | 0.358 | 4.636 | 0.000 | 0.000 |
| Tremateia sp_1 | 1.981 | -1.484 | 0.331 | -4.476 | 0.000 | 0.000 |
| Neoascochyta desmazieri_1 | 1.981 | 1.499 | 0.337 | 4.455 | 0.000 | 0.000 |
| Sporobolomyces sp_2 | 5.117 | -1.711 | 0.405 | -4.223 | 0.000 | 0.000 |
| Epicoccum sp_3 | 1.919 | -1.421 | 0.338 | -4.203 | 0.000 | 0.000 |
| Bulleromyces albus_2 | 1.874 | 1.390 | 0.334 | 4.156 | 0.000 | 0.000 |
| Stagonosporopsis valerianellae_1 | 1.839 | -1.336 | 0.335 | -3.986 | 0.000 | 0.001 |
| Didymella sp_7 | 1.785 | 1.291 | 0.329 | 3.928 | 0.000 | 0.001 |
| Pleosporales_2 | 1.751 | 1.286 | 0.331 | 3.886 | 0.000 | 0.001 |
| Colletotrichum cliviae_1 | 75.491 | 2.274 | 0.592 | 3.843 | 0.000 | 0.001 |
| Didymella sp_2 | 1.732 | 1.229 | 0.325 | 3.781 | 0.000 | 0.001 |
| Epicoccum sp_4 | 1.669 | 1.152 | 0.320 | 3.596 | 0.000 | 0.003 |
| Cladosporium ramotenellum_49 | 1.626 | -1.083 | 0.311 | -3.487 | 0.000 | 0.004 |
| Keissleriella poagena_2 | 1.617 | 1.084 | 0.316 | 3.428 | 0.001 | 0.005 |
| Didymella sp_8 | 1.625 | -1.083 | 0.318 | -3.406 | 0.001 | 0.005 |
| Murispora sp_2 | 1.635 | -1.095 | 0.323 | -3.390 | 0.001 | 0.005 |
| Keissleriella sp_1 | 1.625 | -1.083 | 0.320 | -3.383 | 0.001 | 0.005 |
| Stagonospora sp_1 | 32.898 | -1.793 | 0.530 | -3.380 | 0.001 | 0.005 |
| Cladosporium ramotenellum_15 | 1.617 | -1.072 | 0.317 | -3.379 | 0.001 | 0.005 |
| Cladosporium ramotenellum_13 | 1.617 | -1.072 | 0.317 | -3.378 | 0.001 | 0.005 |
| Alternariaster sp_2 | 1.599 | 1.061 | 0.315 | 3.369 | 0.001 | 0.005 |
| Cladosporium ramotenellum_5 | 1.599 | 1.061 | 0.317 | 3.347 | 0.001 | 0.005 |
| Didymellaceae_2 | 74.438 | 1.912 | 0.604 | 3.166 | 0.002 | 0.009 |
| Bulleromyces albus_1 | 63.929 | 1.652 | 0.546 | 3.025 | 0.002 | 0.013 |
| Phaeosphaeria lunariae_1 | 8.243 | -1.291 | 0.452 | -2.857 | 0.004 | 0.020 |
| Blakeslea trispora_1 | 7.247 | -1.089 | 0.427 | -2.552 | 0.011 | 0.047 |
| Didymellaceae_3 | 66.125 | -1.324 | 0.575 | -2.304 | 0.021 | 0.082 |
| Fusarium sp_1 | 61.919 | 1.209 | 0.590 | 2.048 | 0.041 | 0.128 |

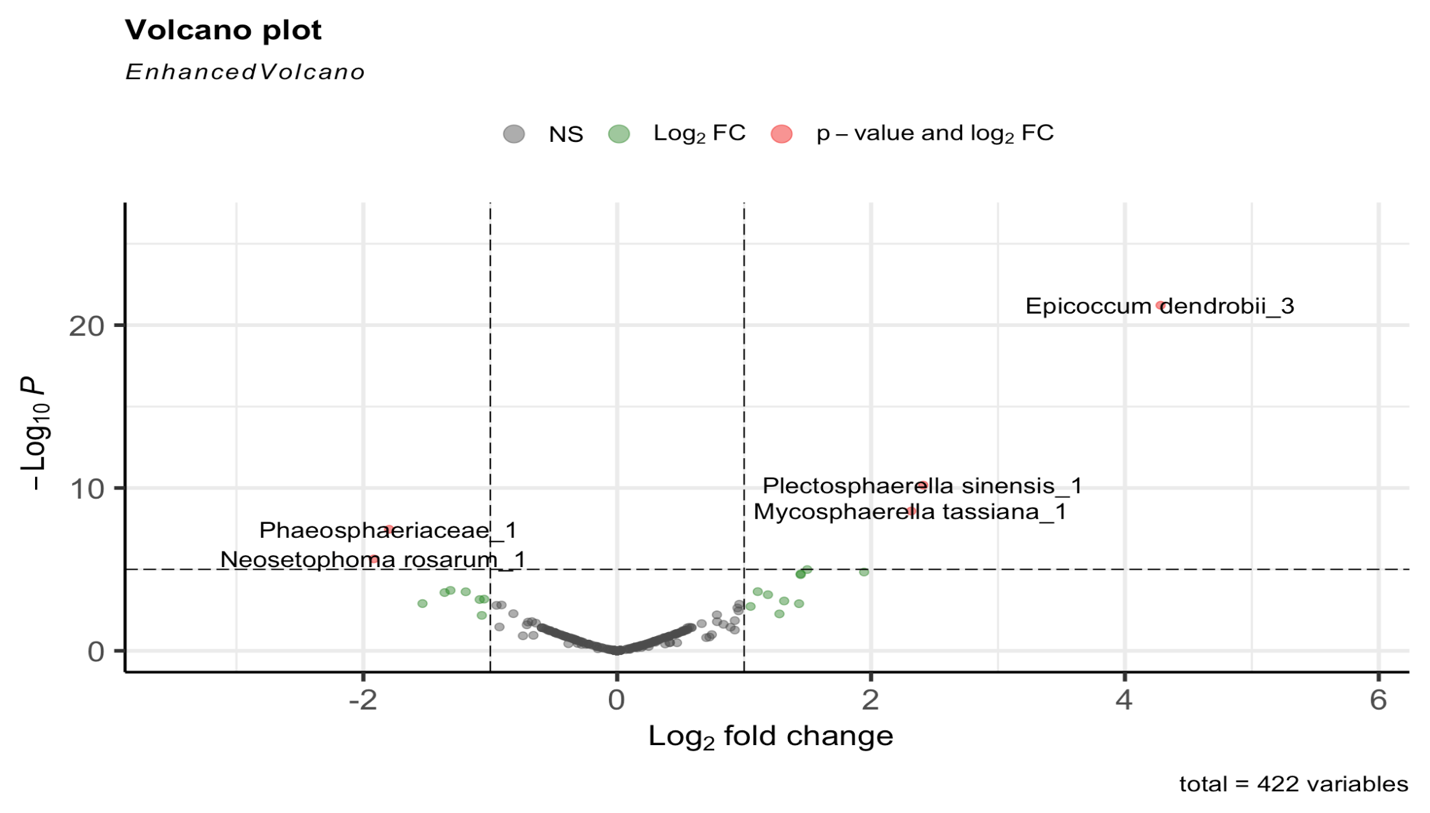

Figure S6. Differential abundance shows log fold change for fungal ASVs detected with the ITS1 primer set. ASVs with positive values to the right of 0 on the x-axis are found in greater abundance in the elevated CO_2_ plots whereas ASVs to the left are found in greater abundance in the control (ambient CO_2_) plots.

Table S2. Fungal taxa identified with the ITS1 primer set with statistically significant log fold change. List has been further subset to those ASVs with log fold change scores >1. Positive log fold change scores indicate higher abundance in the elevated CO_2_ plots whereas negative scores indicate higher abundance in the control/ambient CO_2_ plots.

|  | Base Mean | log2  FoldChange | lfcSE | stat | pvalue | padj |
| --- | --- | --- | --- | --- | --- | --- |
| Epicoccum dendrobii_3 | 10.540 | 4.281 | 0.445 | 9.630 | 0.000 | 0.000 |
| Plectosphaerella sinensis_1 | 3.818 | 2.411 | 0.369 | 6.528 | 0.000 | 0.000 |
| Mycosphaerella tassiana_1 | 4.503 | 2.316 | 0.389 | 5.954 | 0.000 | 0.000 |
| Phaeosphaeriaceae_1 | 2.335 | -1.797 | 0.326 | -5.518 | 0.000 | 0.000 |
| Neosetophoma rosarum_1 | 5.977 | -1.918 | 0.405 | -4.729 | 0.000 | 0.000 |
| Hannaella zeae_1 | 2.548 | 1.497 | 0.339 | 4.412 | 0.000 | 0.000 |
| Albifimbria verrucaria_1 | 11.512 | 1.945 | 0.448 | 4.337 | 0.000 | 0.000 |
| Epicoccum dendrobii_4 | 1.927 | 1.445 | 0.338 | 4.280 | 0.000 | 0.000 |
| Epicoccum dendrobii_9 | 1.927 | 1.445 | 0.340 | 4.253 | 0.000 | 0.000 |
| Sordariomycetes_1 | 3.259 | -1.314 | 0.353 | -3.727 | 0.000 | 0.002 |
| Neosetophoma rosigena_1 | 1.634 | 1.107 | 0.301 | 3.681 | 0.000 | 0.002 |
| Keissleriella taminensis_1 | 1.713 | -1.193 | 0.325 | -3.676 | 0.000 | 0.002 |
| Phaeosphaeria ampeli_1 | 3.721 | -1.360 | 0.373 | -3.648 | 0.000 | 0.002 |
| Colletotrichum spaethianum_1 | 2.157 | 1.187 | 0.333 | 3.570 | 0.000 | 0.002 |
| Neosetophoma rosarum_2 | 1.599 | -1.048 | 0.308 | -3.401 | 0.001 | 0.004 |
| Epicoccum dendrobii_7 | 1.626 | -1.084 | 0.320 | -3.386 | 0.001 | 0.004 |
| Fusarium sporotrichioides_1 | 4.315 | 1.315 | 0.395 | 3.330 | 0.001 | 0.005 |
| Paraboeremia sp_1 | 10.932 | -1.534 | 0.475 | -3.228 | 0.001 | 0.006 |
| Pseudopithomyces rosae_1 | 79.579 | 1.432 | 0.444 | 3.223 | 0.001 | 0.006 |
| Gibberella intricans_3 | 2.185 | 1.051 | 0.338 | 3.107 | 0.002 | 0.008 |
| Bulleromyces albus_1 | 30.941 | 1.278 | 0.460 | 2.781 | 0.005 | 0.019 |
| Septoria cretae_1 | 4.225 | -1.067 | 0.393 | -2.714 | 0.007 | 0.022 |

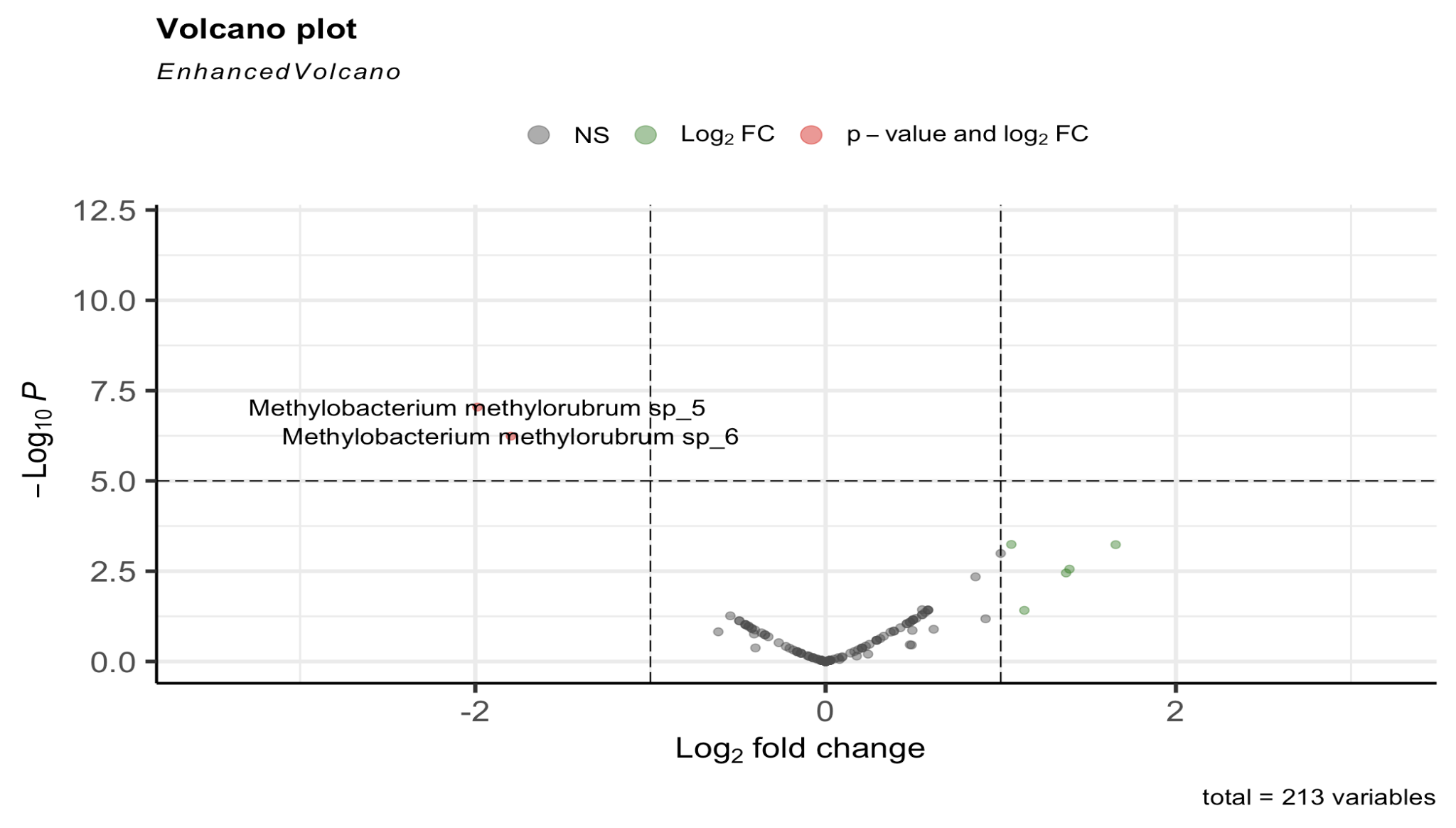

Figure S7. Differential abundance shows log fold change for bacterial ASVs detected with the 16S primer set. ASVs with positive values to the right of 0 on the x-axis are found in greater abundance in the elevated CO_2_ plots whereas ASVs to the left are found in greater abundance in the control/ambient CO_2_ plots.

Table S3. Bacterial taxa identified with the 16s primer set with statistically significant log fold change. List has been further subset to those ASVs with log fold change scores >1. Positive log fold change scores indicate higher abundance in the elevated CO_2_ plots whereas negative scores indicate higher abundance in the control/ambient CO_2_ plots.

|  | Base Mean | log2 Fold Change | lfcSE | stat | pvalue | padj |
| --- | --- | --- | --- | --- | --- | --- |
| Methylobacterium methylorubrum sp_5 | 2.585 | -1.988 | 0.372 | -5.345 | 0.000 | 0.000 |
| Methylobacterium methylorubrum sp_6 | 2.328 | -1.797 | 0.359 | -4.999 | 0.000 | 0.000 |
| Sphingomonas sp_5 | 1.593 | 1.060 | 0.308 | 3.444 | 0.001 | 0.031 |
| Pseudokineococcus lusitanus_1 | 28.988 | 1.656 | 0.481 | 3.440 | 0.001 | 0.031 |
| Aurantimonas sp_1 | 12.488 | 1.392 | 0.465 | 2.992 | 0.003 | 0.098 |
| Sphingomonas sp_1 | 27.895 | 1.373 | 0.471 | 2.917 | 0.004 | 0.108 |
| Escherichia shigella sp_1 | 69.485 | 1.135 | 0.548 | 2.072 | 0.038 | 0.628 |

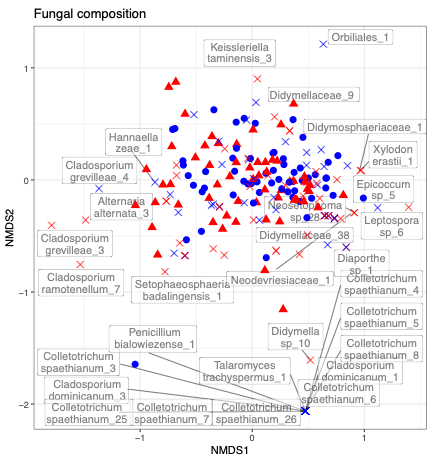

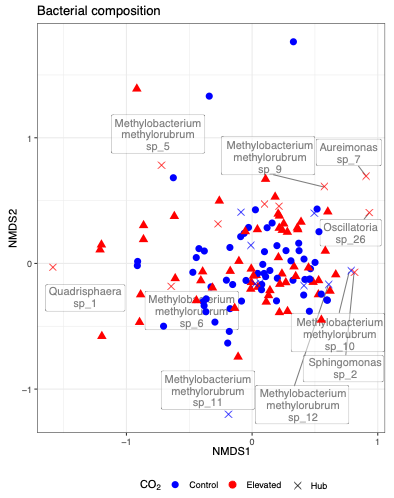

Figure S8. **A.** Soybean foliar fungal composition differed between elevated (red triangles) and ambient (blue circles) CO_2_ treatments (PERMANOVA, R^2^= 0.031, p =0.001). Hub taxa that had influential NMDS scores are also labeled (for the fungal dataset hubs with NMDS scores larger than the absolute value of 0.8 are displayed). Shown are community composition along NMDS1 and NMDS2 based on the ITS2 primer set, and community composition results were largely analogous for the ITS1 primer set. **B.** Soybean foliar bacterial composition based on the 16S primer set did not differ between elevated and ambient [CO_2_] treatments (PERMANOVA, R^2^= 0.012, p =0.16). Hub taxa hubs with NMDS scores larger than the absolute value of 0.5 are displayed as the bacterial dataset has fewer identified hubs.
